# PyTorch-FEA: Autograd-enabled Finite Element Analysis Methods with Applications for Biomechanical Analysis of Human Aorta

**DOI:** 10.1101/2023.03.27.533816

**Authors:** Liang Liang, Minliang Liu, John Elefteriades, Wei Sun

## Abstract

**Motivation:** Finite-element analysis (FEA) is widely used as a standard tool for stress and deformation analysis of solid structures, including human tissues and organs. For instance, FEA can be applied at a patient-specific level to assist in medical diagnosis and treatment planning, such as risk assessment of thoracic aortic aneurysm rupture/dissection. These FEA-based biomechanical assessments often involve both forward and inverse mechanics problems. Current commercial FEA software packages (e.g., Abaqus) and inverse methods exhibit performance issues in either accuracy or speed.

**Methods:** In this study, we propose and develop a new library of FEA code and methods, named PyTorch-FEA, by taking advantage of autograd, an automatic differentiation mechanism in PyTorch. We develop a class of PyTorch-FEA functionalities to solve forward and inverse problems with improved loss functions, and we demonstrate the capability of PyTorch-FEA in a series of applications related to human aorta biomechanics. In one of the inverse methods, we combine PyTorch-FEA with deep neural networks (DNNs) to further improve performance.

**Results:** We applied PyTorch-FEA in four fundamental applications for biomechanical analysis of human aorta. In the forward analysis, PyTorch-FEA achieved a significant reduction in computational time without compromising accuracy compared with Abaqus, a commercial FEA package. Compared to other inverse methods, inverse analysis with PyTorch-FEA achieves better performance in either accuracy or speed, or both if combined with DNNs.

## 1. Introduction

Structural Finite-element analysis (FEA) has been widely used to perform stress and deformation analysis of complex structures, for which an analytical solution might be infeasible. In the biomedical domain, FEA has been applied to study the mechanics of human tissues and organs, tissue-medical device interactions, diagnostic and treatment strategies at a patient-specific level [1–11], which may need to solve forward and inverse mechanics problems. For instance, an emerging application of FEA for medical diagnosis is the risk assessment of aortic aneurysms. Aortic aneurysm ranks consistently in the top 20 causes of death in the U.S. population [12]. Thoracic aortic aneurysm (TAA) is manifested as an abnormal bulging of thoracic aortic wall, and it is a leading cause of death in adults with a prevalence of ~ 1% in the general population [13, 14]. From the perspective of biomechanics, TAA rupture/dissection occurs when the stress acting on the aortic wall exceeds the material strength of the wall [15]. To accurately obtain the stress distribution of TAA, FEA is often used as the core solution tool in the risk assessment workflow [10, 11, 16–19]. This workflow typically involves both forward (compute stress given stress-free geometry, boundary conditions, and material properties) and inverse (identify material properties given loaded geometries and boundary conditions) problems. Commercial FEA software packages (e.g., Abaqus) have been used for aortic wall stress analysis of human aorta [10, 11, 19]. The forward problem can be solved by these FEA packages with graphical interfaces. However, inverse problem solvers are not supported by these FEA packages and typically require an optimization framework that calls a FEA package to run FEA simulations iteratively to update parameters. These existing inverse strategies [20–24] heavily rely on FEA-updating that treats FEA as a blackbox, which results in well-known performance issues in either accuracy or speed.

In this study, we propose and develop PyTorch-FEA, a new library of FEA code and methods to solve forward and inverse problems with significantly improved performance. PyTorch [25] is an open source machine learning library, especially for developing deep neural networks (DNNs), and it can run on CPU and GPU. Besides those specific to DNNs, it has a high performance linear algebra module and many extensions [26], which are suitable for FEA implementation. The key benefit from PyTorch is the automatic differentiation mechanism, named autograd, which implements backpropagation from the loss, the fundamental algorithm to train DNNs. It enables the design of new loss functions for solving inverse problems using FEA principles, and the optimization can be done in a backpropagation fashion. It eliminates the need for hand-calculating derivatives (e.g., the stiffness matrix), which simplifies the implementation of FEA methods. Naturally, it enables a seamless integration of FEA and DNNs, as will be shown in this study.

We demonstrate the advantages of PyTorch-FEA in four basic applications for biomechanical analysis of human aorta, which are: (1) forward simulation of aorta inflation from zero-pressure to a target pressure, given a hyperelastic constitutive model with known material parameters; (2) inverse estimation of ex-vivo material parameters, given the zero-pressure geometry and a pressurized geometry; (3) inverse estimation of the zero-pressure geometry, given a pressurized geometry and the material parameters; and (4) inverse estimation of in-vivo material parameters, given two pressurized geometries at two pressure levels. To evaluate the performance in each application, we employ FEA-generated data for which “ground-truth” solutions are known.

The application-1 is the basic forward analysis to obtain the deformation and stress of the aorta at a target blood pressure, e.g., 18kPa representing the systolic pressure at hypertension stage-1 [27]; and if the stress is high, the aortic aneurysm rupture risk is also high. In this application, the initial undeformed geometry, blood pressure, and material parameters are given as the inputs, and the analysis outputs are the deformed geometry and stress at the target blood pressure. We compare PyTorch-FEA with the commercial software Abaqus and show that the discrepancy is negligible and PyTorch-FEA is much faster.

The application-2 arises from the study of human aorta material properties in inflation experiments [28]. In such an experiment, an aortic root, which is obtained from a cadaver heart, is inflated from 0 to ~26kPa by smoothly injecting saline solution to the root that is placed inside a container; markers are placed on the root surface to enable camera-based marker tracking; and then pressure-strain responses are obtained from the experiment data. Subsequently, tissue material parameters of a specific hyperelastic constitutive model can be determined by fitting the pressure-strain responses in an inverse analysis. In this application, the initial (i.e., undeformed, zero-pressure) geometry, the deformed geometry at a known blood pressure, and the form of the constitutive model are given as the inputs, and the inverse analysis estimates the material parameters of the constitutive model.

The application-3 arises from the fact that the geometry of the aorta in clinical images is in the pressurized state and therefore cannot be used as the initial, undeformed geometry in a forward FEA (e.g., application-1) to obtain stresses; and therefore the zero-pressure (i.e., undeformed) geometry needs to be estimated by using a deformed geometry at a known blood pressure and a specific constitutive model with known material parameters. This application also arises from the potential use of 3D-printed tissue-engineered aortic root as a replacement of the native aortic root [29]. In such case, the engineered aortic root is load-free and needs to match the deformed geometry of the native aortic root when pressurized. In this application, we will show two PyTorch-FEA inverse methods, and one of the methods combines FEA with DNNs to speed up the inverse analysis process while maintaining the same accuracy.

In the application-4, the material parameters of the aortic wall tissue are considered unknown and will be estimated by using two pressurized geometries at two different pressure levels, assuming the form of the constitutive model is given. In practice, the two geometries can be reconstructed from multiphase 3D CT images of a patient’s aorta [24], one from the diastolic phase and the other from the systolic phase. By using patient-specific material parameters in a forward FEA, more accurate stress distribution can be obtained, which may improve the accuracy of aortic aneurysm diagnostics. In this application, we will demonstrate the capability of PyTorch-FEA using statical determinacy to solve this inverse problem, and we will use two different loss functions to show that the combination of residual force term and stress error term leads to better accuracy.

We will open-source Pytorch-FEA on GitHub when the paper is published.

## 2. Methods

### 2.1 Material modeling of human aortic wall tissue

Soft biological tissues, such as human aortic wall tissue, comprise bundles of collagen fibers embedded in a ground matrix and can be regarded as fiber-reinforced composites, which exhibit nonlinear hyperelastic behaviors [15–18]. Modeling of the mechanical behavior (a.k.a. constitutive modeling) of hyperelastic tissues is often achieved by specifying the strain energy Ψ per unit undeformed volume as a function of strain invariants and fiber orientations. These strain invariants are derived from the deformation gradient tensor ***F***.

Among existing hyperelastic constitutive relations, the GOH model [30] is widely used for the aortic wall tissue. In this model, tissues are composed of a matrix material with two families of embedded fibers, each of which has a preferred direction. The strain energy density function is expressed by:

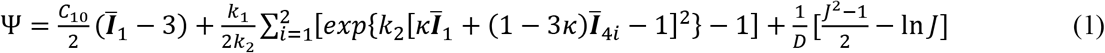

The parameter *C*_10_ describes the matrix material. The parameters *k*_1_ and *k*_2_ describe the fiber properties. The deviatoric strain invariants 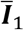 and 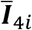 characterize the deformation of the matrix and preferred fiber directions, respectively. The parameter *κ* describes the distribution of the fiber orientation. The parameter *θ* defines the mean local fiber direction in a local coordinate system. *D* is a constant to enforce material incompressibility. *J* = det (***F***).

Using a constitutive model, different stresses can be calculated, including the first PK stress tensor ***P*** and the Cauchy stress tensor ***σ***

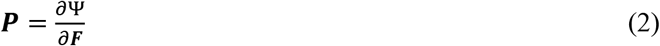

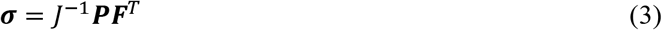

In this work, we use the GOH model to demonstrate the capabilities of PyTorch-FEA. We select a representative set of the material parameters, *C*_10_ = 54.73, *k*_1_ = 2225.62, *k*_2_ = 24.61, *κ* = 0.2494, *θ* = 32.43 from our previous studies [10, 11, 19, 31].

### 2.2 Finite element formulation in the context of solid mechanics

We provide a brief overview of the finite element formulation, and we try to keep the notations and descriptions to be as consistent as possible with those in the book [32]. The finite element formulation can be established by using the virtual work representation of the equilibrium equation, and the directional derivative of the total potential energy Π yields the principle of virtue work, which is given by

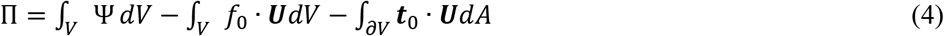

The body force *f*_0_ is ignored for the applications of human aorta. ***U*** is the displacement of a material point from the initial position ***X*** to the current position ***x***, i.e., ***x*** = ***X*** + ***U**. V* represents the undeformed body of the object. ***t***_0_ is traction force on the surface *∂V* of the object. The total potential energy Π can also be expressed using integrals evaluated on the deformed body. Given the constitutive model, the external forces, and the geometry of the object at the initial state, the total potential energy will decrease and approach a minimum at equilibrium.

In the FEA formulation, the domain of interest is discretized into finite elements. The position ***x*** of a material point inside an element with *n_e_* nodes can be interpolated using the node positions {***x**_i_*, *i* = 1,…, *n_e_*} and a shape function *N_i_*:

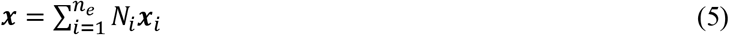

The same interpolation can be applied to the initial position ***X*** and the displacement ***U*** of the same material point.

As a result, the deformation gradient tensor at the material point located at ***x*** inside an element can be calculated by

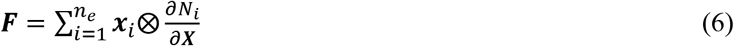

By setting the directional derivative of the total potential energy to zero (equivalent to the virtual work principal), the internal equivalent nodal force 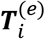 and the external equivalent nodal force 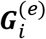 can be obtained [32]:

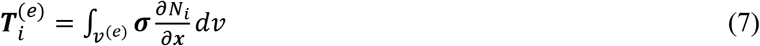

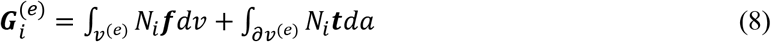

In the above equation, *ν*^(*e*)^ represents an element of the deformed body and its index is *e*; ***t*** is traction force acting on the surface of the element; and ***f*** is body force that is ignored for the applications of human aorta. The integrals are evaluated numerically using the Gaussian integration method.

A node may be shared by adjacent elements, and therefore, at each node, the forces from neighbor elements are added together to obtain the assembled internal equivalent nodal force *T_i_* and the assembled external equivalent nodal force *G_i_* at the node-*i* :

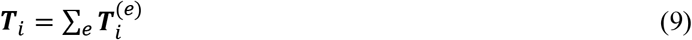

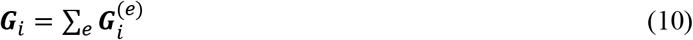

At the equilibrium state, the nodal residual force ***R**_i_* is zero:

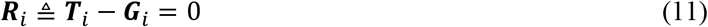

The relationship among Π, ***R**_i_*, and the displacement ***U**_i_* of the node-*i* is given by

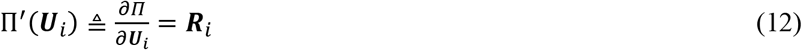

Commercial FEA software packages (e.g., Abaqus) implement forward FEA solvers to determine the deformed geometry, given the initial (i.e., undeformed) geometry, external loading, boundary conditions, and a specific constitutive model with known parameters. A known variable in the forward problem becomes unknown (i.e., to be solved) in an inverse problem. Solution of inverse problems requires substantial changes to the FEA implementation, which is often not feasible in commercial FEA software packages. The applications 2-4 mentioned in the introduction section are inverse problems and are readily tractable by functionalities in Pytorch-FEA.

### 2.3 The design of PyTorch-FEA

The 3-level structure of PyTorch-FEA is shown in Figure 1. On the element level, it has basic functions to perform interpolation using shape functions (Eq. (5)), calculate the deformation gradient tensor (Eq.(6)), perform numerical integration to compute the equivalent nodal forces (Eq.(7) and Eq.(8)) and the strain energy stored in an element (i.e., *∫_V_*^(*e*)^ Ψ *dV*). We implemented the hexahedral element that is similar to the C3D8 element in Abaqus. To avoid the volumetric locking issue associated with nearly incompressible materials (e.g., Eq.(1)), we implemented selective-reduced integration [33] for the numerical integrations. In addition to the hexahedral element, we also implemented functions to handle blood pressure acting on the inner surface of the aortic wall, and functions for surface integrals are implemented for calculating the equivalent pressure (i.e., external) force at each node of a quadrilateral element. It also defines the interfaces to constitutive models, which must provide three functions: one to calculate the strain energy density (e.g., Eq.(1)) at a material point (e.g., integration point) given deformation gradient tensor ***F*** and element orientation as inputs, and the other two to calculate the stress tensors ***P*** and ***σ***. We note that PyTorch provides automatic differentiation (a.k.a. autograd) for almost all operations, thus, the stress tensor ***P*** can be computed directly by autograd using Eq.(2). These element-level functions are implemented in batch-mode to handle all of the elements at the same time.

**Figure 1.**
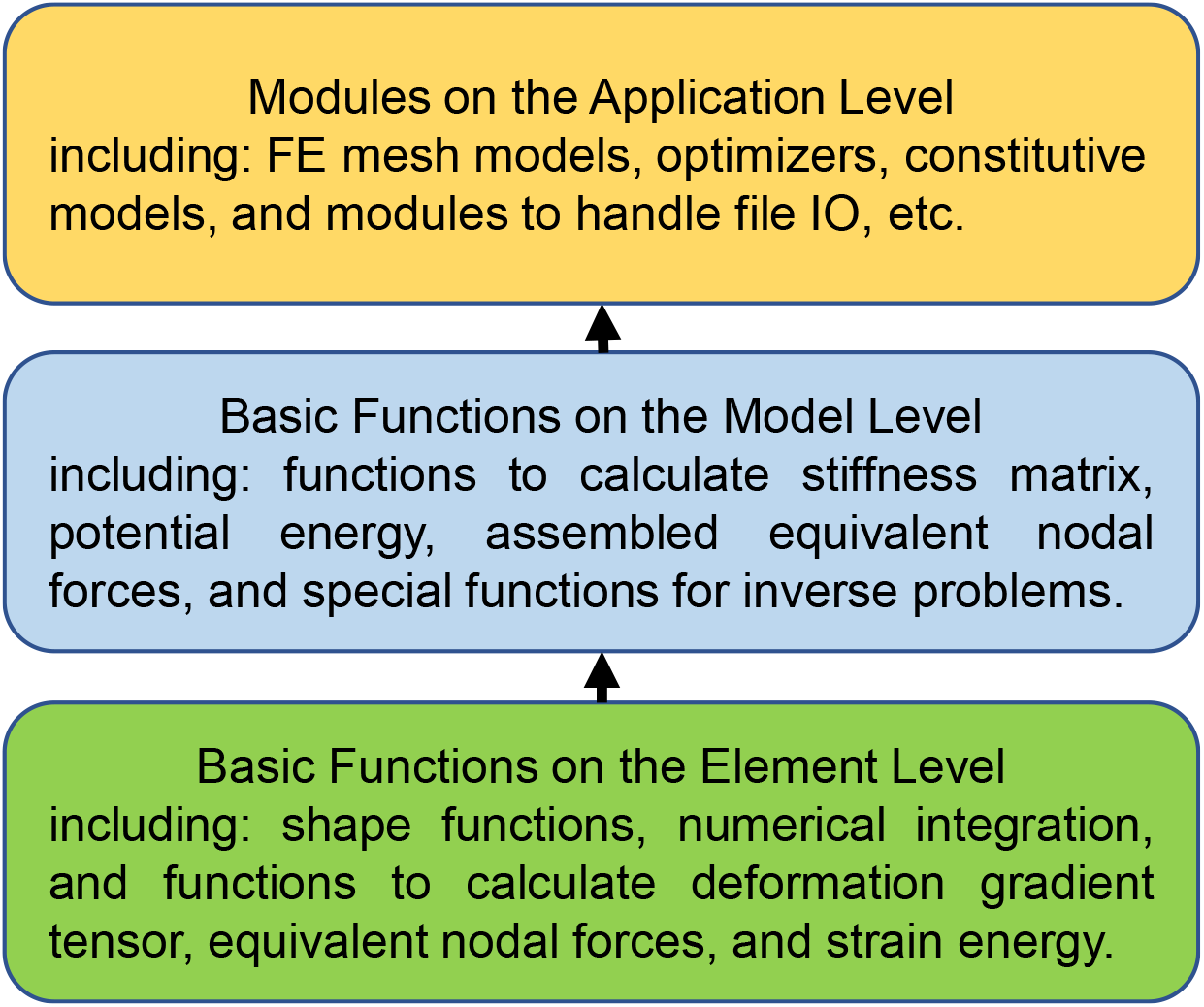
The three-level design of Pytorch-FEA

On the model level, PyTorch-FEA has basic functions to calculate the total potential energy (Eq.(4)), assemble the equivalent nodal forces (Eq.(9) and Eq.(10)) at each node, and calculate the stiffness matrix that is useful for optimization. The stiffness matrix is the second-order derivative of the total potential energy with respect to the displacement, which is also called Hessian matrix. And therefore, it is the first-order derivative of the residual force with respect to the displacement. The size of the full stiffness matrix is 3*N*×3*N*, where *N* is the total number of nodes. The stiffness matrix is very sparse. A naive implementation would be directly using PyTorch autograd to compute the derivative and obtain a dense matrix that will consume lots of CPU or GPU memory, leading to OOM (out-of-memory) error. Instead, we provide a memory-efficient implementation to assemble the sparse matrix from individual elements.

On the application level, PyTorch-FEA currently only supports the four applications of aorta biomechanical analysis. It has FE models of the aortic wall, which are implemented as Python class, encapsulating all necessary functions to calculate a variety of quantities (e.g., stress, deformation, strain energy, etc). The inner surface needs to be specified, i.e., the quad elements on which the pressure load will be applied. Currently, it only uses one type of boundary condition: zero displacement at each inlet/outlet boundary of the aorta. It also includes optimizers and constitutive models, as well as modules to handle file IO. The four applications are explained in the next sections.

### 2.4 The application of aorta inflation simulation

As shown in Figure 2, in this application, the initial (i.e., undeformed) geometry, blood pressure, and the constitutive model with known parameters (Section 2.1) are given as the inputs, and the objective is to obtain the deformed geometry at the specified blood pressure and then calculate stress. Here, we consider blood pressure distribution on the inner wall to be uniform, although Pytorch-FEA supports non-uniform distributions. The blood pressure value is set to 18kPa, representing the systolic pressure at hypertension stage-1 [27]. If the stress at the hypertension stage-1 is high, the aneurysm rupture risk is also high.

**Figure 2.**
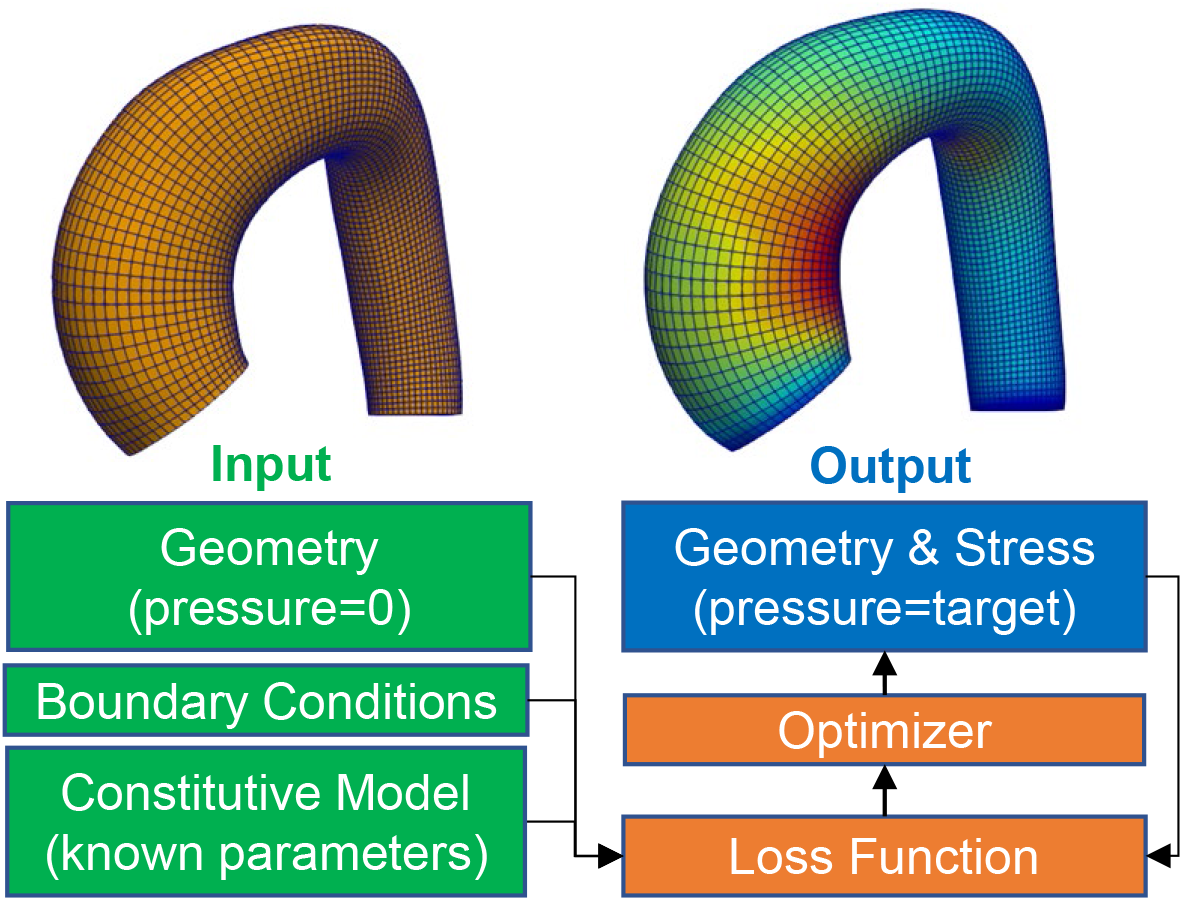
Forward simulation of aorta inflation

We use the total potential energy in Eq.(4) as a pseudo loss function of the displacement field **U** of the nodes. We note that Eq.(4) is not used to calculated the residual force by autograd; and instead, the residual force is calculated by using Eq.(11). For optimization, we implemented a quasi-newton method based on the L-BFGS optimizer [34]. Briefly, the Taylor expansion could be applied to the total potential energy:

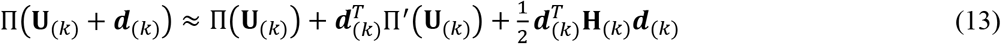

In the above equation, **U**_(κ)_ represents the displacement field (assembled into a vector) of all the free nodes at the current iteration. ***d***_(*κ*)_ is a small increment. **H**_(*κ*)_ is known as the tangent stiffness matrix at the current iteration-*k*, and 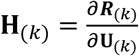, where ***R***_(*κ*)_ is the residual force field of the nodes at the current iteration. If the boundary nodes are fixed with zero-displacement, the corresponding rows and columns in **H**_(*κ*)_ need to be removed. The optimal increment ***d***_(*κ*)_, which minimizes the quadratic form (i.e., the right-side of Eq.(13)), is known as

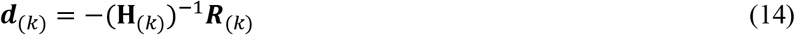

In practice, the inverse (**H**_(*κ*))_^-1^ is never calculated, and the following sparse linear system of equations is solved to obtain ***d***_(*κ*)_:

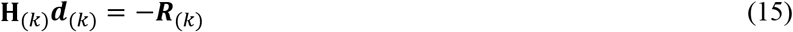

We used a Python package PyPardiso [35] as the sparse solver. Also, a line search is applied to adjust the increment:

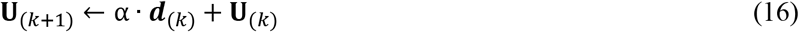

where the nonnegative scalar α is auto-adjusted to minimize the loss function.

A full-Newton method will solve Eq.(15) in every iteration, and therefore it will be very computationally expensive and time-consuming. The L-BFGS method [34] avoids Eq.(15) by approximating the Hessian inverse, and therefore each iteration is fast, but it may need numerous iterations to converge due to the imprecise approximation. As a combination of the two methods, we re-initialize L-BFGS with Eq.(15) every 20 iterations, and this combined method works well in this application.

We use the following metric as the primary metric to determine the convergence of optimization:

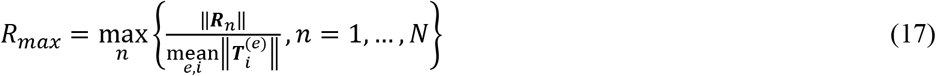

We use ‖■‖ to denote scalar absolute value, vector L2 norm, or matrix Frobenius norm in this paper. ***R**_n_* is the residual force (Eq.(11)) at the node-*n* at the current iteration, and the total number of nodes of an aorta mesh is *N*. If *R_max_ < threshold*, then the optimization is considered converged. In this application, the threshold is 0.005. According to Abaqus documentation, a similar metric is used by Abaqus FEA solver to determine convergence. We also use two secondary metrics to monitor the optimization process: ΔU*_ratio_* = ‖**U**_(*κ*+1)_ – **U**_(*κ*)_‖/‖**U**_(*κ*+1)_‖ and 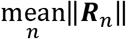 at the current iteration. In our application, ΔU*_ratio_* is negligibly small when *R_max_* < 0.005.

To improve convergence, we use pseudo time steps similar to those in Abaqus. The simulation duration is mapped to a time interval between 0 and 1, and the interval is divided into small steps. The external load is increased gradually by *pressure*_(*t*)_ = *t* × *pressure* where *pressure* is the target pressure (e.g., 18kPa), and *t* is the current time in the interval. For each time point *t*, the optimizer runs until *R_max_* < *threshold* to obtain the optimal displacement under *pressure*_(*t*)_. If it does not converge at the current time point, the program will go back to the previous time point with a randomly-chosen time step. If it converges, then the time step will increase but not exceed the pre-defined max step (e.g., 0.01 in this application).

To demonstrate the capability of PyTorch-FEA for this application, we use seven representative initial geometries sampled from a statistical shape model (SSM) [19]. The SSM is built on 60 real aorta geometries in our previous studies [31], and it captures the major shape variations. Among the seven geometries, six of those are the mean shape ± 1.5 × mode (mode 1, 2 or 3) of shape variations, and the other one is the mean shape. Figures of the initial (i.e., undeformed, zero-pressure) geometries are provided in the appendix. The results are in Section 3, which show that the discrepancy between PyTorch-FEA and Abaqus is negligible.

### 2.5 The application of ex-vivo material parameter estimation

In this application, the initial (i.e., undeformed, zero-pressure) geometry, the deformed geometry at a known blood pressure, and the form of the constitutive model are given, and the objective is to estimate the material parameters of the constitutive model. We use the same seven test cases in Section 2.4 to demonstrate this application, and the deformed geometries are obtained by the PyTorch-FEA forward method.

To achieve the objective, we define the loss function to be:

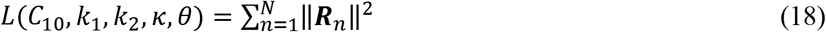

We use L-BFGS with line search as the optimizer. The optimization converged in a few seconds for each test case, and the difference between the estimated parameters and the true parameters is negligible. Because of the automatic differentiation capability of Pytorch-FEA, the gradient of the loss with respect to each parameter can be exactly calculated (e.g., 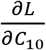), and therefore, we can use gradient-based optimization that runs much faster than non-gradient-based optimization that is typically used in FEA-updating inverse methods (e.g., [20–22]) that may use the following loss function:

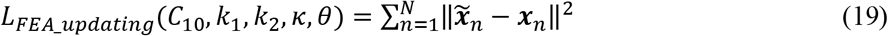

where 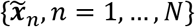 is an estimation of the deformed-geometry obtained by running a forward FEA with the current estimations of the material parameters. Because *L_FEA_updating_* is not differentiable with respect to the material parameters, the FEA-updating inverse method needs numerous forward FEA simulations.

### 2.6 The application of zero-pressure geometry estimation

In this application, the zero-pressure (i.e., initial, undeformed) geometry needs to be estimated by using the information about the deformed geometry at a known blood pressure and the constitutive model with known parameters. To demonstrate this application, we use the same seven test cases in Section 2.5 and set the pressure to be 10kPa, representing in vivo diastolic pressure.

In a previous study [19], we used the backward-displacement method [36] to obtain the zero-pressure geometry of human aorta, and it was implemented by using Abaqus, Python, and Matlab. The key iteration step of the method is given by:

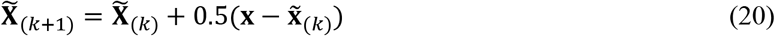

where 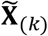 represents the estimated zero-pressure geometry at the iteration *k*, 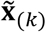 represents the deformed-geometry obtained by a forward FEA using 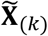 as the initial geometry, and **x** represents the true deformed-geometry that is deformed from the true but unknown zero-pressure geometry. At the initialization step, 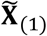 is equal to **x**. The total number of iterations is fixed to 20, leading to long computation time based on our previous study [19]. We note that *k* is the index of an iteration (i.e., a forward FEA simulation), not the index of a node. Although this backward-displacement method is easily implemented into PyTorch-FEA, we employ two better inverse strategies in Pytorch-FEA.

To take advantage of the automatic differentiation capability of Pytorch-FEA, we propose two new methods: PyFEA-P0 and PyFEA-NN-P0, where P0 stands for pressure zero, NN stands for neural network, and PyFEA refers to Pytorch-FEA. The procedure of PyFEA-P0 is very similar to the forward FEA of the application-1 in Section 2.4, and the displacement field is the primary variable that will be optimized, and then

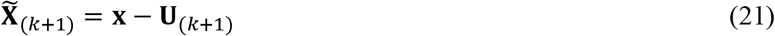

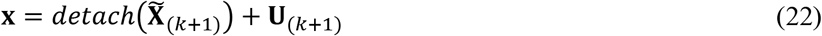

The Eq.(22) does nothing mathematically, but it is necessary to obtain the correct derivatives related to the displacement. The autograd mechanism works with a computational graph that is automatically constructed and represents a sequence of operations to compute the loss. By detaching 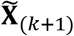 from the computational graph, Eq.(22) will ensure the correct derivatives as if a forward FEA is performed starting from 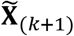. Similarly, the element orientation (a function of 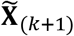 variable needs to be detached from the graph. To improve convergence, we use pseudo time steps counting down from 1 to 0.

The second method PyFEA-NN-P0 needs an autoencoder DNN that can be explained by two functions: an encoder function: ***c*** = *f_encoder_*({***U**_n_*, *n* = 1,2, … *N*}) and a decoder function: {***U**_n_*, *n* = 1,2,… *N*} = *f_decoder_* (***c***). Here, ***U**_n_* is the displacement of the node-*n* from the zero-pressure state to the pressurized state, and *N* is the total number of nodes of an aorta mesh. Briefly, the input to the encoder is the entire displacement field of an aorta mesh, and the displacement field is compressed into a lower dimensional code vector ***c*** . Given the code vector as input, the decoder recovers the displacement field. In this application, the encoder is implemented by an MLP (multi-layer perceptron) that has two hidden layers with 128 units per layer and Softplus activations, and the dimensions of code vector ***c*** is 16; and the decoder is implemented by another MLP mirroring the structure of the encoder MLP. The autoencoder is trained on 273 samples generated by the SSM (Section 2.4), which do not include the seven test cases. After training, the decoder will be used in the optimization process, and the code vector ***c*** becomes the primary variable to be optimized. For each of the test cases, the optimal value of *c* should lead to the minimum of the loss (i.e., total potential energy). The L-BFGS optimizer with line search is used, and the gradient can be calculated by

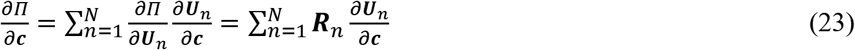

where 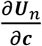 is the derivative of the decoder output with respect to its input and is auto-calculated by autograd.

### 2.7 The application of in-vivo material parameter estimation

In this application, the material parameters of the aortic wall tissue are considered unknown and need to be estimated by using two pressurized geometries at two different pressure levels, and the form of the constitutive model is given by Eq.(1). In practice, the two geometries can be reconstructed from gated 3D CT scans of a patient’s aorta [24], at two cardiac phases (diastolic and systolic). The identified patient-specific material parameters can be used in forward FEA to calculate patient-specific wall stress under elevated blood pressure, which can lead to more accurate risk assessment of TAA [31, 37]. To demonstrate the capability of PyTorch-FEA for this application, we use the same seven test cases in Section 2.4, and for each test case, the two pressurized geometries are obtained by the PyTorch-FEA forward method using pressure values of 10kPa (diastole) and 16kPa (systole).

To achieve the objective, we employ the following loss function by requiring diminishing residual force and matching stresses according to the principle of statical determinacy [38–40]:

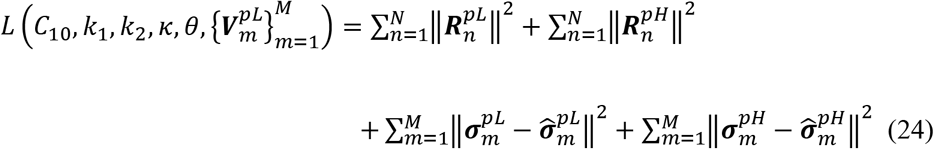

In the loss function, 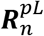 is the residual force at the node-*n* at the diastolic phase (pL stands for pressure low), and 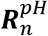 is the residual force at the node-*n* at the systolic phase (pH stands for pressure high). 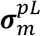 and 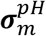 are stress tensors at the element-*m*, which are computed by using Eq.(3) at the diastolic phase and the systolic phase, respectively. *N* is the number of nodes, and *M* is the number of elements of an aorta mesh. 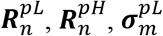, and 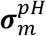 are nonlinear functions of the unknown material parameters {*C*_10_, *k*_1_, *k*_2_, *κ*, 0}. The stress tensors 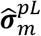 and 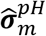 are directly obtained using the principle of statical determinacy [38–40] with the two pressurized geometries and are independent of the material parameters. 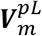 is the left stretch tensor at the element-*m* from the zero-pressure state to the diastolic phase, and it is unknown. It is related to the deformation gradient tensor 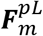 by the polar-decomposition: 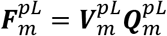, where 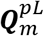 is rotation. We use the L-BFGS method with line search as the optimizer for this application.

From the two pressurized geometries, the deformation gradient tensor 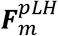 of the element-*m* from the diastolic phase to the systolic phase (pLH stands for pressure from low to high) can be directly calculated. Then, the deformation gradient tensor 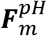 of the element-*m* from the zero-pressure state to the systolic phase can be obtained by 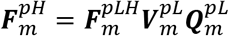. Although 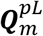 is unknown, it will not affect stress calculations using the constitutive model, and the explanation is provided in the appendix. Thus, the stress tensors (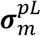 and 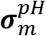) and the residual forces (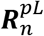 and 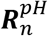) can be calculated by using the current values/estimations of the deformation gradient tensors and the current estimations of the material parameters. The boundary nodes and elements are removed from the loss function because 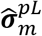 and 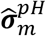 are inaccurate near the boundaries due to the known boundary effect.

To obtain 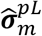 and 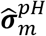 using the principle of statical determinacy for each test case, we first apply the inverse method PyFEA-P0 (Section 2.6) to recover the zero-pressure geometries, for which the material parameters are set to *C*_10_ = 10000, *k*_1_ = 0, *k*_2_ = 1, *κ* = 0, *θ* = 0, representing a very stiff material. Then, the zero-pressure geometries are inflated by the PyTorch-FEA forward method (Section 2.4) to the diastolic phase and the systolic phase to obtain the stresses 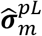 and 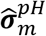, respectively.

If we remove the residual force terms, i.e., 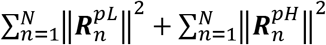, from the loss function, the loss function will become similar to the loss in our previous method [41] that has a very complex implementation using co-rotational coordinate system for problem formulation and using Abaqus, Python, and Matlab for programming. In the result section, we will show that the residual force term improves the estimation accuracy.

## 3. Results

The results of the application-1 in Section 2.4 are reported in Table 1. The differences between Pytorch-FEA and Abaqus for forward simulations in this application are quantified for each test case by using three metrics related to geometry and stress:

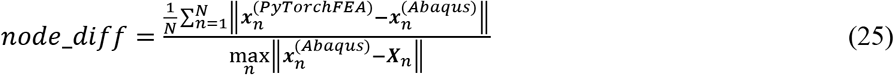

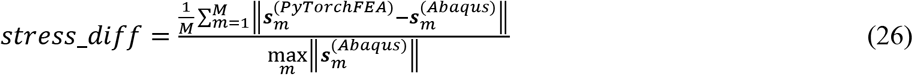

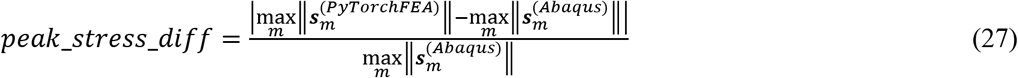

We use von Mises stress as 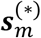 in Eq.(26) and Eq.(27) because the stresses in Abaqus are always in the local coordinate systems of the elements in the deformed geometry, and the stresses in Pytoch-FEA are always in the global coordinate system. Pytorch-FEA and Abaqus were tested on the same computer equipped with an Intel i7-8700 CPU and two Nvidia Titan V GPUs. Abaqus only uses the CPU with FP64 data type (i.e., double-precision). PyTorch-FEA uses the CPU and one GPU with FP64 data type. The sparse solver in Pytorch-FEA runs on CPU and consumes about 1/3 of the total time cost. For each test case, the time cost of Pytorch-FEA is about 65% of the time cost of Abaqus.

**Table 1:** The differences between PyTorch-FEA and Abaqus in the application-1 in Section 2.4

| Test Case Index | *node_diff* | *stress_diff* | *peak_stress_diff* |
| --- | --- | --- | --- |
| 1 | 0.00107 | 0.00197 | 0.00988 |
| 2 | 0.00081 | 0.00172 | 0.00264 |
| 3 | 0.00044 | 0.00164 | 0.01510 |
| 4 | 0.00125 | 0.00119 | 0.00041 |
| 5 | 0.00194 | 0.00201 | 0.00412 |
| 6 | 0.00232 | 0.00187 | 0.00114 |
| 7 | 0.00176 | 0.00146 | 0.00108 |

The results of the application-2 in Section 2.5 are reported in Table 2. The error of each estimated material parameter is quantified by the following metric:

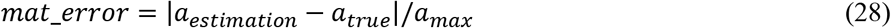

where *a* represents a parameter and *a_max_* is the upper bound of the parameter. The lower bound of each parameter is close to zero. According to our previous studies [31, 37], *C*_10_max_ = 120 (kPa), *k*_1_max_ = 6000 (kPa), *k*_2_max_ = 60, *κ_max_* = 1/3, and *θ_max_* = 90 (degree). For each test case, the time cost is about 10 seconds.

**Table 2:**
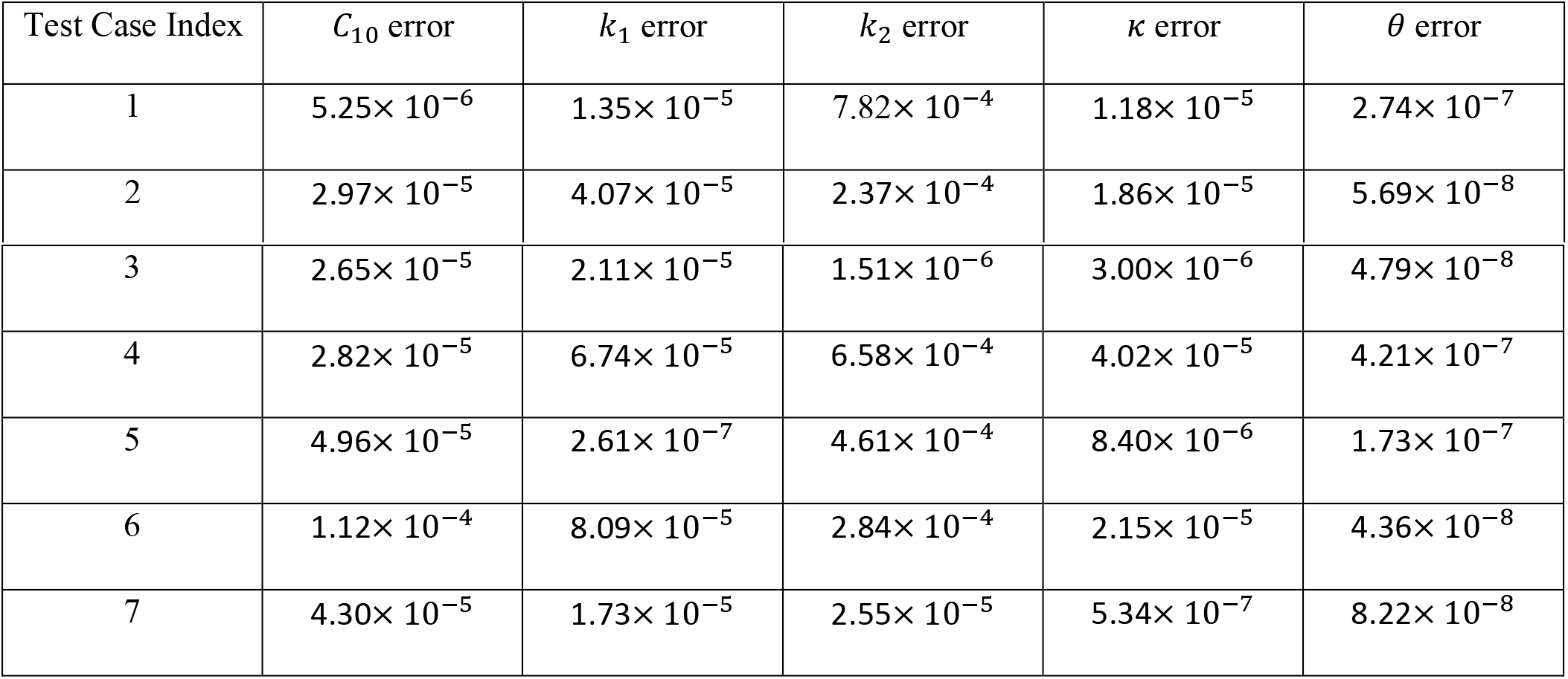
The errors of the estimated material parameters in the application-2 in Section 2.5

The results of the application-3 in Section 2.6 are reported in Table 3, comparing the performance of the three inverse methods, including backward-displacement (BD), PyFEA-p0, and PyFEA-NN-p0. For each test case, we use the following metrics to measure the performance of a method:

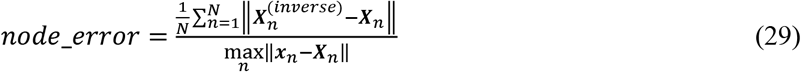

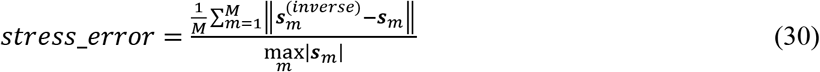

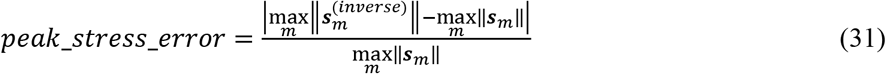

***x**_n_* is the position of the node-*n* of the aorta mesh model at the diastolic phase. 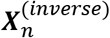 is the estimated position of the node-*n* at the zero-pressure state. ***s**_m_* is the true stress at the element-*m* at the diastolic phase, and 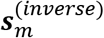 is the estimated stress from an inverse method. To be consistent with the evaluation in application-1, we use von Mises stress in Eq.(30) and Eq.(31). To compare the methods, we calculate the average of each metric (averaged across the seven cases) as showing in Table 3. The performance of each individual method for the seven test cases is reported in the Appendix. The fastest method is PyFEA-NN-P0 that only takes less than 10 seconds to handle a test case. The slowest method is PyFEA-P0 that may take hours for a test case. The speed of the BD method is between the other two, but its accuracy is relatively lower.

**Table 3:** Performance comparison of the three inverse methods in the application-3 in Section 2.6

| Method | average *node_error* | average *stress_error* | average *peak_stress_error* |
| --- | --- | --- | --- |
| PyFEA-NN-P0 | 0.00190 | 0.00035 | 0.00066 |
| PyFEA-P0 | 0.01254 | 0.00026 | 0.00070 |
| BD | 0.08942 | 0.00334 | 0.00911 |

The results of the application-4 in Section 2.7 are reported in Table 4, comparing the performance of the two inverse methods, one with the residual force terms in the loss (Method-A, the second row in Table 4), and the other using only the stress terms in the loss (Method-B, the third row in Table 4). The metric of Eq.(28) is also used in this application. The performance of each individual method for the seven test cases is reported in the Appendix. PyFEA-p0 is used in this application, and it only takes a few minutes to handle a test case. The difference between the estimated stresses (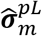 and 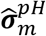) and the true stresses is about 3%, and the details are reported in the Appendix. With residual force terms in the loss function, Method-A has lower errors in *C*_10_, *k*_2_, *κ*, and *θ*. The two methods have roughly the same time cost.

**Table 4:** Performance comparison of the two inverse methods in the application-4 in Section 2.7

| Method | $C_{10}$ error<br>(average) | $k_1$ error<br>(average) | $k_2$ error<br>(average) | $\kappa$ error<br>(average) | $\theta$ error<br>(average) |
| --- | --- | --- | --- | --- | --- |
| with residual<br>force terms | 0.03921 | 0.02140 | 0.03015 | 0.01755 | 0.00551 |
| without residual<br>force terms | 0.11146 | 0.01825 | 0.03438 | 0.01836 | 0.00840 |

## 4. Discussion

The capability of PyTorch-FEA has been demonstrated on the four applications. The application-1 shows that PyTorch-FEA forward analysis solution is almost the same as Abaqus, and the small discrepancy (< 0.5% in stress as shown in Table 1) could be caused by the differences in the optimizers. PyTorch-FEA eliminates the need for manual-calculating derivatives (e.g., calculating stiffness matrix, calculating stress by differentiating the strain energy density function, etc), and therefore the implementation is clean and efficient for GPU. The current implementation of the PyTorch-FEA forward method is about 1.5 times faster than Abaqus, and the speed is limited by the sparse solver that runs on CPU and accounts for 1/3 of the time cost. We will upgrade the sparse solver with a GPU implementation [42] in our future work to further reduce the time cost. Also, if provided with the current generation of GPUs, e.g., Nvidia H100, which is magnitudes faster than Titan V GPU used in this study, PyTorch-FEA can become much faster. The applications 2-4 show the advantage of PyTorch-FEA for solving the inverse problems. For each inverse problem, we only need to define an appropriate loss function, and the optimization is similar to training a DNN by backpropagation from the loss, which enables us to define new or improved loss functions (Eq.(18), Eq.(23), and Eq.(24)) to achieve better accuracy or faster speed.

PyTorch-FEA enables a natural marriage of FEA and DNNs, which enjoys the benefits of both: accurate and fast, as shown in the application-3 for inverse estimation of the zero-pressure geometry. In our inverse method PyFEA-NN-P0, the DNN decoder is a generative model that can generate a displacement field from an input code vector. Then, the solution space is reduced from the dimension of 30000 (1000 nodes in a mesh model, and each node has three displacement values in 3D) to a much lower dimension of 16. As a result, the optimization can complete in seconds, magnitudes faster than the other methods. If a much larger training dataset is available, the decoder could be replaced by a generative adversarial network (GAN) [43] or a diffusion model [44], which are state-of-the-art generative models for image and natural language data. This FEA-DNN integration could be potentially used for the other applications, to further speed up the analysis.

Recently, we and other research groups have developed constitutive models based on DNNs [45–51], which perform better than the expert-prescribed models in some applications. However, it can be difficult to implement a complex DNN as a user subroutine in the programming language that is compatible with commercial software packages, e.g., FORTRAN for Abaqus UMAT. PyTorch-FEA could potentially serve as a platform for developing and testing DNN-based constitutive models.

Physics-informed neural networks (PINNs) [52], which attracted a lot of attentions in the field of scientific machine learning, offers a direct solution to PDEs by using DNNs and physics principles. Although PINNs are intriguing, current PINNs have not led to performance improvement compared with commercial FEA methods in the solid mechanics domain and suffer from the error/instability [53] in the deformation gradient tensor calculated by differentiating a DNN with respect to the input: 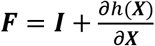 where *h*(***X***) is a DNN to output displacement. The instability is not a numerical defect of autograd, and it is closely related to the well-known issue of DNN robustness [54], and it is known that [55, 56] gradients of a DNN can change dramatically if the input changes only a little or even if the same DNN is trained twice on the same data.

Several open-source FEA software packages/programs are available, such as FEBio [57], FEAP [58], and FEniCS [59], which employ traditional Newton-Rapson forward FEA solvers. FEAP and FEniCS are general-purpose FEA programs. FEBio specializes in solving nonlinear large deformation (forward) problems in biomechanics and biophysics, and it works for the application-1 but does not have functionalities for the inverse applications. The official FEniCS code does not have off-the-shelf functionalities to support complex hyperelastic constitutive models with selective-reduced integration, geometry, loading and boundary condition in this study. A recent work enhanced FEniCS capability for a compressible hyperelastic model [60] to solve inverse problems using the FEA updating method (Eq. (19)). FEBio and FEAP do not support autograd. There has been an attempt to combine FEniCS and PyTorch to solve simple PDEs [61], but it is unclear if it could be extended to complex cases, such as the applications in this study.

The current PyTorch-FEA only supports the four applications for biomechanical analysis of the human aorta. The basic functions on the element level and the model level are general FEA functions, not specifically designed for the four applications, which enable extensions of PyTorch-FEA. We will continue developing PyTorch-FEA for applications in the field of biomechanics, especially cardiovascular biomechanics.

## 5. Conclusion

We have presented PyTorch-FEA, a new library of FEA code and methods, representing a new approach to develop FEA methods to forward and inverse problems in solid mechanics. We have demonstrated its capabilities in the four basic applications in aorta biomechanics. PyTorch-FEA eases the development of new inverse methods and enables a natural integration of FEA and DNNs, which will have numerous potential applications.

**Figure 3.**
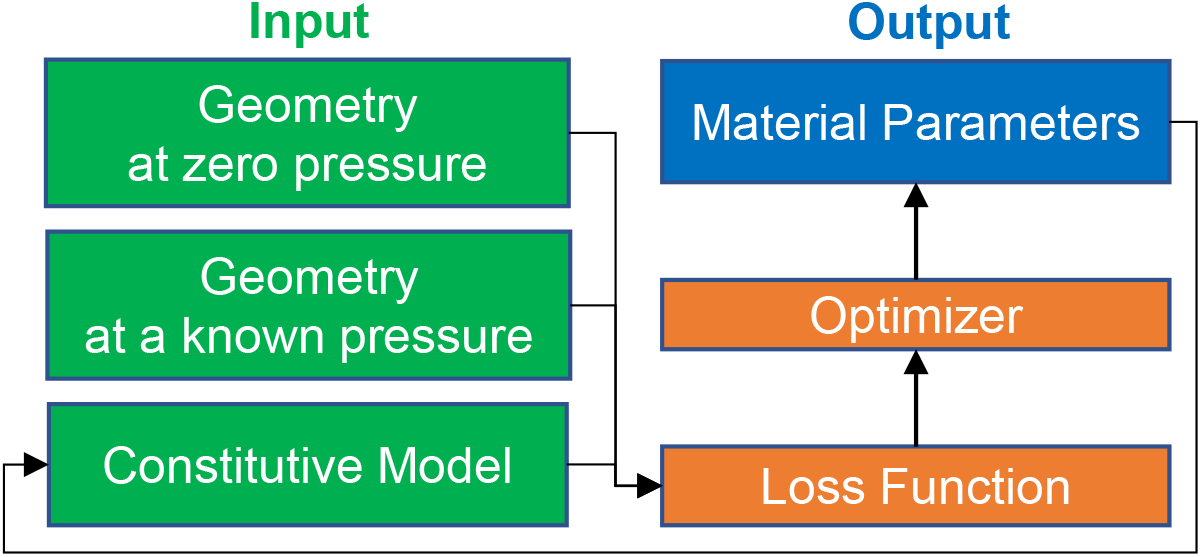
Inverse estimation of ex-vivo material parameters

**Figure 4.**
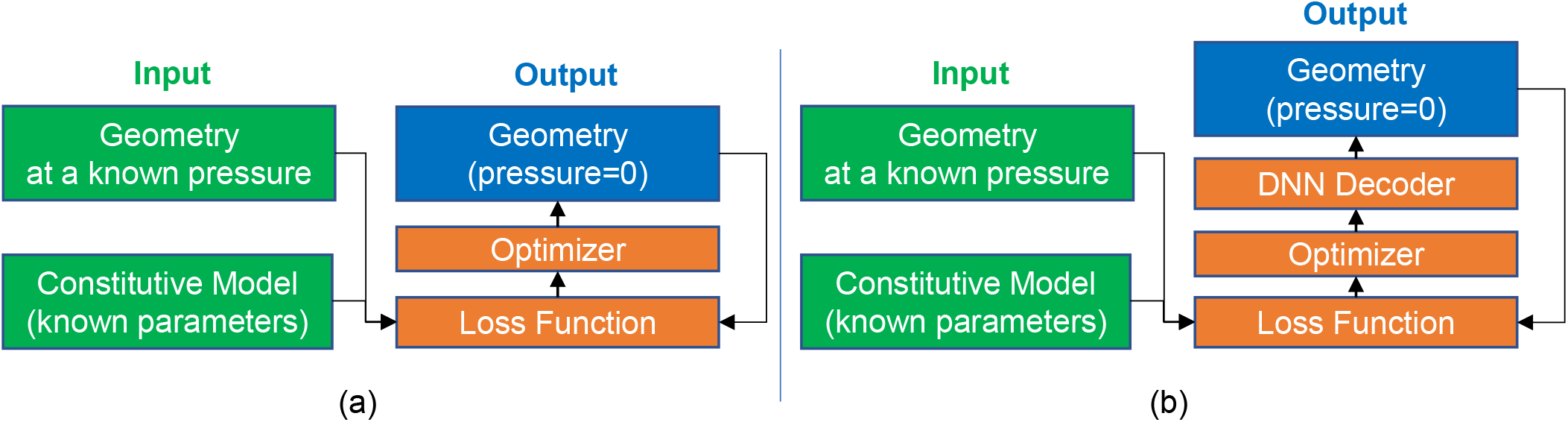
Two PyTorch-FEA inverse methods for the estimation of zero-pressure geometry. (a) The method PyFEA-P0. (b) The method PyFEA-NN-P0 that has a DNN decoder to generate a displacement field.

**Figure 5.**
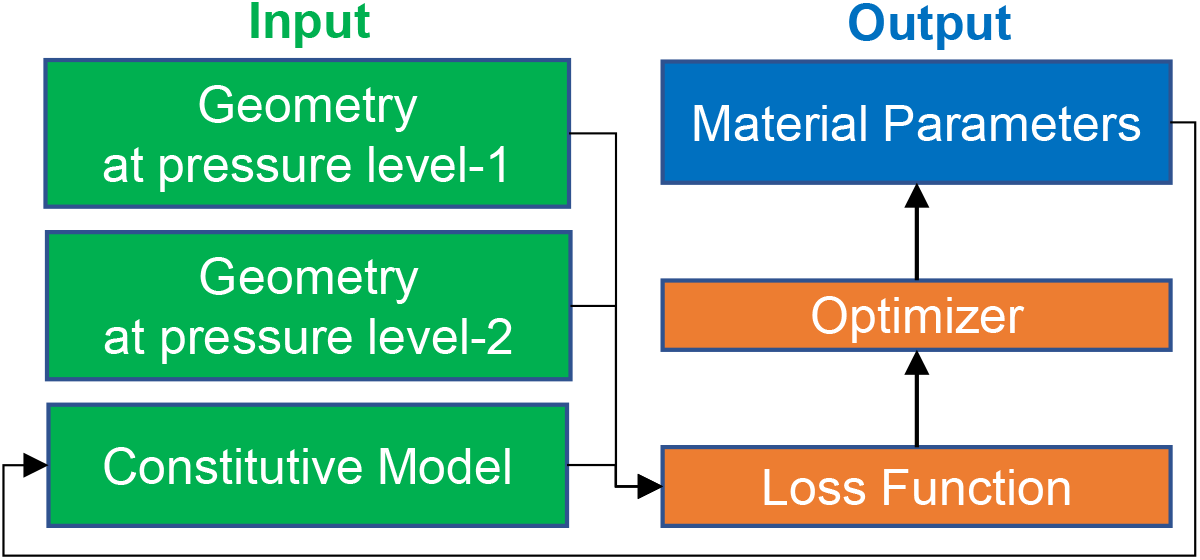
Inverse estimation of in-vivo material parameters

## Acknowledgement

This work was supported in part by the NIH grant R01HL158829.

## Appendix

### 1. The zero-pressure geometries of the seven test cases

**Figure A.1.**
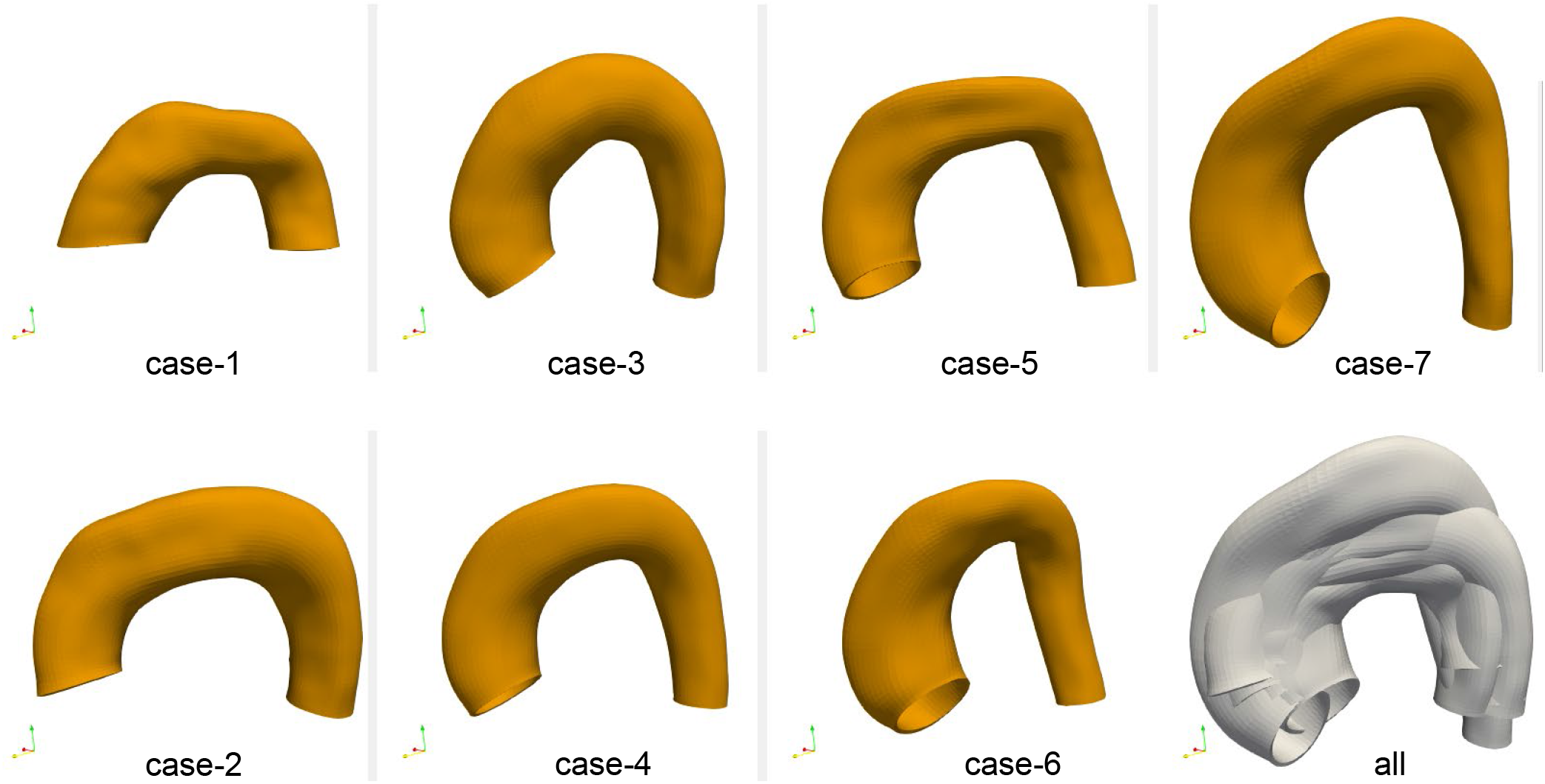
The zero-pressure geometries of the seven test cases. Case-4 is the mean shape.

### 2. Explanation of 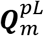 in the application-4 in Section 2.7

In many models (e.g., Eq.(1)), the strain energy density Ψ is an explicit function of the strain invariants of the right Cauchy–Green tensor ***C*** ≜ ***F**^T^ **F***, and the strain invariants are independent of any rotation. We can calculate 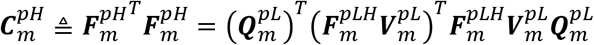, and therefore the strain invariants of 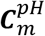 are independent of the rotation 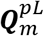. Similarly, the strain invariants of 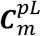 are independent of the rotation 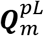.

The GOH model in Eq.(1)has terms related to fiber orientation ***a*** (a vector in the global coordinate system). 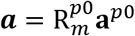, where 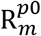 is the coordinate transform matrix (a.k.a. orientation of the element-*m*) from a local coordinate system to the global coordinate system. **a**^*p*0^ is the fiber orientation in the local coordinate system of the zero-pressure geometry, and it is determined by the material parameter *θ*. The GOH model needs to calculate 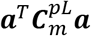 and 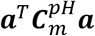 along with several other strain invariants of 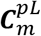 and 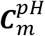. By using the polar decomposition, we have 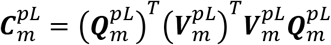, and 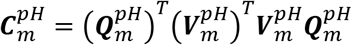. Thus, we obtain 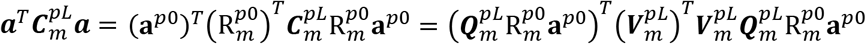. Let 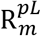 denote the element orientation at the low-pressure sate, and then we have 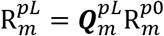 in theory (it may have numerical errors) because 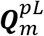 is the rotation from the zero-pressure state to the low-pressure state. Thus, 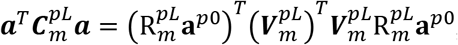, which is independent of 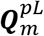. Similarly, it can be shown that 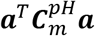 is independent of 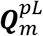.

In summary 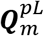 does not affect stress calculations using the constitutive model. 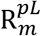 is used as the element orientation input for the constitutive model.

### 3. The individual performance of the three inverse methods in application-3 in Section 2.6

**Table A1.** Performance of PyFEA-NN-p0 in the application-3

| Test Case Index | *node_error* | *stress_error* | *peak_stress_error* |
| --- | --- | --- | --- |
| 1 | 0.001603687 | 0.000505738 | 0.000738402 |
| 2 | 0.001609170 | 0.000380529 | 0.000390757 |
| 3 | 0.001657492 | 0.000350096 | 0.001384317 |
| 4 | 0.001324879 | 0.000219505 | 0.000480732 |
| 5 | 0.002291675 | 0.000376495 | 0.000533053 |
| 6 | 0.002071800 | 0.000311183 | 0.000127871 |
| 7 | 0.002709812 | 0.000312835 | 0.000984382 |

**Table A2.** Performance of PyFEA-p0 in the application-3

| Test Case Index | *node_error* | *stress_error* | *peak_stress_error* |
| --- | --- | --- | --- |
| 1 | 0.013546776 | 0.000472815 | 0.001112776 |
| 2 | 0.010208852 | 0.000231789 | 0.000156242 |
| 3 | 0.008520643 | 0.000258453 | 0.001786788 |
| 4 | 0.007296290 | 0.000188554 | 0.000659276 |
| 5 | 0.026151690 | 0.000238295 | 0.000080900 |
| 6 | 0.005821294 | 0.000222569 | 0.000575342 |
| 7 | 0.016261040 | 0.000236795 | 0.000507866 |

**Table A3.** Performance of Backward-Displacement in the application-3

| Test Case Index | *node_error* | *stress_error* | *peak_stress_error* |
| --- | --- | --- | --- |
| 1 | 0.050443336 | 0.002912068 | 0.007059155 |
| 2 | 0.099223304 | 0.004954056 | 0.015746499 |
| 3 | 0.097816046 | 0.004365298 | 0.028571723 |
| 4 | 0.088351709 | 0.002286753 | 0.005556760 |
| 5 | 0.093116105 | 0.002741628 | 0.005139562 |
| 6 | 0.096268331 | 0.002496273 | 0.001162921 |
| 7 | 0.100715101 | 0.003628388 | 0.000532503 |

### 4. The individual performance of the two inverse methods in application-4 in Section 2.7

**Table A4:** The performance of the Method-A in the application-2 in Section 2.5

| Test Case Index | $C_{10}$ error | $k_1$ error | $k_2$ error | $\kappa$ error | $\theta$ error |
| --- | --- | --- | --- | --- | --- |
| 1 | 0.129247378 | 0.042951700 | 0.064761514 | 0.044543974 | 0.019333531 |
| 2 | 0.000622965 | 0.022907529 | 0.051915732 | 0.026669237 | 0.009406890 |
| 3 | 0.006252636 | 0.005289165 | 0.024329692 | 0.000735397 | 0.000075500 |
| 4 | 0.055838874 | 0.011756801 | 0.006615650 | 0.008653229 | 0.001687726 |
| 5 | 0.015083297 | 0.023531461 | 0.022397862 | 0.014278869 | 0.001958761 |
| 6 | 0.025283741 | 0.017563589 | 0.016656474 | 0.010606607 | 0.000387365 |
| 7 | 0.042121716 | 0.025813757 | 0.024350831 | 0.017392986 | 0.005729018 |

**Table A5:** The performance of the Method-B in the application-2 in Section 2.5

| Test Case Index | $C_{10}$ error | $k_1$ error | $k_2$ error | $\kappa$ error | $\theta$ error |
| --- | --- | --- | --- | --- | --- |
| 1 | 0.192870881 | 0.057150165 | 0.126684566 | 0.054823763 | 0.021151645 |
| 2 | 0.180261673 | 0.036096861 | 0.043758695 | 0.039467320 | 0.019296666 |
| 3 | 0.118783674 | 0.008901363 | 0.011829653 | 0.008886206 | 0.003399421 |
| 4 | 0.138038347 | 0.009141652 | 0.023494923 | 0.002153123 | 0.000827182 |
| 5 | 0.082047669 | 0.003260920 | 0.021085351 | 0.008920925 | 0.005473181 |
| 6 | 0.029286580 | 0.011930520 | 0.004169498 | 0.012040075 | 0.006770422 |
| 7 | 0.038954431 | 0.001235340 | 0.009627799 | 0.002200260 | 0.001858200 |

**Table A6:** The accuracy of 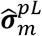 and 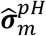 in application-4 in Section 2.7

| Test Case Index | $\hat{\sigma}_m^{pL}$ error | $\hat{\sigma}_m^{pH}$ error |
| --- | --- | --- |
| 1 | 0.042568215 | 0.042068338 |
| 2 | 0.034829138 | 0.032056521 |
| 3 | 0.030086746 | 0.028548364 |
| 4 | 0.023311289 | 0.022395104 |
| 5 | 0.024368868 | 0.022454050 |
| 6 | 0.022995795 | 0.020644430 |
| 7 | 0.019870676 | 0.018354943 |

Note: the error is calculated using the metric in Eq.(26)

The estimation errors shown in Tables A4 and A5 are caused by the errors in 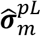 and 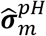.

## Reference

[1] F. Auricchio, M. Conti, S. Morganti, and A. Reali, “Simulation of transcatheter aortic valve implantation: a patient-specific finite element approach,” Comput Methods Biomech Biomed Engin, vol. 17, no. 12, pp. 1347–57, 2014, doi: 10.1080/10255842.2012.746676.

[2] C. Capelli et al., “Patient-specific simulations of transcatheter aortic valve stent implantation,” Med Biol Eng Comput, vol. 50, no. 2, pp. 183–92, Feb 2012, doi: 10.1007/s11517-012-0864-1.

[3] P. de Jaegere et al., “Patient-Specific Computer Modeling to Predict Aortic Regurgitation After Transcatheter Aortic Valve Replacement,” JACC Cardiovasc Interv, vol. 9, no. 5, pp. 508–12, Mar 14 2016, doi: 10.1016/j.jcin.2016.01.003.

[4] H. A. Dwyer, P. B. Matthews, A. Azadani, L. Ge, T. S. Guy, and E. E. Tseng, “Migration forces of transcatheter aortic valves in patients with noncalcific aortic insufficiency,” J Thorac Cardiovasc Surg, vol. 138, no. 5, pp. 1227–33, Nov 2009, doi: 10.1016/j.jtcvs.2009.02.057.

[5] S. Morganti, N. Brambilla, A. S. Petronio, A. Reali, F. Bedogni, and F. Auricchio, “Prediction of patient-specific post-operative outcomes of TAVI procedure: The impact of the positioning strategy on valve performance,” JBiomech, vol. 49, no. 12, pp. 2513–9, Aug 16 2016, doi: 10.1016/j.jbiomech.2015.10.048.

[6] S. Morganti et al., “Simulation of transcatheter aortic valve implantation through patient-specific finite element analysis: two clinical cases,” J Biomech, vol. 47, no. 11, pp. 2547–55, Aug 22 2014, doi: 10.1016/j.jbiomech.2014.06.007.

[7] W. Sun, C. Martin, and T. Pham, “Computational modeling of cardiac valve function and intervention,” Annu Rev Biomed Eng, vol. 16, pp. 53–76, Jul 11 2014, doi: 10.1146/annurev-bioeng-071813-104517.

[8] Q. Wang, S. Kodali, C. Primiano, and W. Sun, “Simulations of transcatheter aortic valve implantation: implications for aortic root rupture,” Biomech Model Mechanobiol, vol. 14, no. 1, pp. 29–38, Jan 2015, doi: 10.1007/s10237-014-0583-7.

[9] Q. Wang, E. Sirois, and W. Sun, “Patient-specific modeling of biomechanical interaction in transcatheter aortic valve deployment,” J Biomech, vol. 45, no. 11, pp. 1965–71, Jul 26 2012, doi: 10.1016/j.jbiomech.2012.05.008.

[10] C. Martin, W. Sun, and J. Elefteriades, “Patient-specific finite element analysis of ascending aorta aneurysms,” American Journal of Physiology-Heart and Circulatory Physiology, vol. 308, no. 10, pp. H1306–H1316, 2015, doi: 10.1152/ajpheart.00908.2014.

[11] C. Martin, W. Sun, T. Pham, and J. Elefteriades, “Predictive biomechanical analysis of ascending aortic aneurysm rupture potential,” Acta Biomaterialia, vol. 9, no. 12, pp. 9392–9400, 2013/12/01/ 2013, doi: https://doi.org/10.1016/j.actbio.2013.07.044.

[12] CDC. “Centers for Disease Control and Prevention, National Center for Injury Prevention and Control, WISQARS Leading Causes of Death Reports, 1999 - 2018: https://webappa.cdc.gov/cgi-bin/broker.exe.“ (accessed Sept 15, 2020).

[13] T. Faggion Vinholo, M. A. Zafar, B. A. Ziganshin, and J. A. Elefteriades, “Nonsyndromic Thoracic Aortic Aneurysms and Dissections—Is Screening Possible?,” Seminars in Thoracic and Cardiovascular Surgery, 2019/06/15/ 2019, doi: https://doi.org/10.1053/j.semtcvs.2019.05.035.

[14] A. Verstraeten, I. Luyckx, and B. Loeys, “Aetiology and management of hereditary aortopathy,” Nature Reviews Cardiology, Review Article vol. 14, p. 197, 01/19/online 2017, doi: 10.1038/nrcardio.2016.211.

[15] S. Sherifova and and G. A. Holzapfel, “Biochemomechanics of the thoracic aorta in health and disease,” Progress in Biomedical Engineering, vol. 2, no. 3, p. 032002, 2020/07/14 2020, doi: 10.1088/2516-1091/ab9a29.

[16] T. C. Gasser, “The Biomechanical Rupture Risk Assessment of Abdominal Aortic Aneurysms—Method and Clinical Relevance,” in Biomedical Technology: Modeling, Experiments and Simulation, P. Wriggers and T. Lenarz Eds. Cham: Springer International Publishing, 2018, pp. 233–253.

[17] T. C. Gasser, “Biomechanical Rupture Risk Assessment: A Consistent and Objective Decision-Making Tool for Abdominal Aortic Aneurysm Patients,” (in eng), Aorta (Stamford), vol. 4, no. 2, pp. 42–60, 2016, doi: 10.12945/j.aorta.2015.15.030.

[18] S. Sherifova and G. A. Holzapfel, “Biomechanics of aortic wall failure with a focus on dissection and aneurysm: A review,” Acta Biomaterialia, vol. 99, pp. 1–17, 2019/11/01/ 2019, doi: https://doi.org/10.1016/j.actbio.2019.08.017.

[19] L. Liang, M. Liu, C. Martin, J. A. Elefteriades, and W. Sun, “A Machine Learning Approach to Investigate the Relationship between Shape Features and Numerically Predicted Risk of Ascending Aortic Aneurysm,” Biomechanics and Modeling in Mechanobiology, vol. 16, no. 5, pp. 1519–1533, 2017.

[20] A. Wittek et al., “In vivo determination of elastic properties of the human aorta based on 4D ultrasound data,” Journal of the Mechanical Behavior of Biomedical Materials, vol. 27, pp. 167–183, 11// 2013, doi: http://dx.doi.org/10.1016/j.jmbbm.2013.03.014.

[21] A. Wittek et al., “A finite element updating approach for identification of the anisotropic hyperelastic properties of normal and diseased aortic walls from 4D ultrasound strain imaging,” Journal of the Mechanical Behavior of Biomedical Materials, vol. 58, pp. 122–138, 5// 2016, doi: http://dx.doi.org/10.1016/j.jmbbm.2015.09.022.

[22] M. Liu, L. Liang, and W. Sun, “Estimation of in vivo mechanical properties of the aortic wall: A multiresolution direct search approach,” Journal of the Mechanical Behavior of Biomedical Materials, vol. 77, pp. 649–659, 2018/01/01/ 2018, doi: https://doi.org/10.1016/j.jmbbm.2017.10.022.

[23] M. Liu et al., “On the Identification of Heterogeneous Nonlinear Material Properties of the Aortic Wall from Clinical Gated CT Scans,” International Conference on Biomechanics and Medical Engineering, 2019.

[24] M. Liu et al., “Identification of in vivo nonlinear anisotropic mechanical properties of ascending thoracic aortic aneurysm from patient-specific CT scans,” Scientific Reports, vol. 9, no. 1, p. 12983, 2019.

[25] A. Paszke et al., “PyTorch: An Imperative Style, High-Performance Deep Learning Library,” Neural Information Processing Systems, pp. 8024–8035, 2019.

[26] M. Fey and J. E. Lenssen, “Fast Graph Representation Learning with PyTorch Geometric,” ICLR Workshop on Representation Learning on Graphs and Manifolds, 2019.

[27] A. Hooker, K. G. Buda, and M. Pasha, “Managing stage 1 hypertension: Consider the risks, stop the progression,” Cleveland Clinic Journal of Medicine, vol. 89, no. 5, p. 244, 2022, doi: 10.3949/ccjm.89a.21101.

[28] K. Li, Q. Wang, T. Pham, and W. Sun, “Quantification of structural compliance of aged human and porcine aortic root tissues,” (in eng), J Biomed Mater Res A, vol. 102, no. 7, pp. 2365–74, Jul 2014, doi: 10.1002/jbm.a.34884.

[29] Z. Qian et al., “Quantitative Prediction of Paravalvular Leak in Transcatheter Aortic Valve Replacement Based on Tissue-Mimicking 3D Printing,” (in eng), JACC Cardiovasc Imaging, vol. 10, no. 7, pp. 719–731, Jul 2017, doi: 10.1016/j.jcmg.2017.04.005.

[30] T. C. Gasser, R. W. Ogden, and G. A. Holzapfel, “Hyperelastic modelling of arterial layers with distributed collagen fibre orientations,” Journal of The Royal Society Interface, vol. 3, no. 6, pp. 15–35, 2005/09/28 2005, doi: 10.1098/rsif.2005.0073.

[31] M. Liu et al., “A probabilistic and anisotropic failure metric for ascending thoracic aortic aneurysm risk assessment,” Journal of the Mechanics and Physics of Solids, vol. 155, p. 104539, 2021/10/01/ 2021, doi: https://doi.org/10.1016/j.jmps.2021.104539.

[32] J. Bonet and R. D. Wood, Nonlinear Continuum Mechanics for Finite Element Analysis. Cambridge University Press, 2008.

[33] S. Doll, K. Schweizerhof, R. Hauptmann, and C. Freischläger, “On volumetric locking of low-order solid and solid-shell elements for finite elastoviscoplastic deformations and selective reduced integration,” Engineering Computations, vol. 17, pp. 874–902, 2000.

[34] D. C. Liu and J. Nocedal, “On the limited memory BFGS method for large scale optimization,” Mathematical Programming, vol. 45, no. 1, pp. 503–528, 1989/08/01 1989, doi: 10.1007/BF01589116.

[35] A. Haas, “PyPardiso,” GitHub repository https://github.com/haasad/PyPardisoProject, 2023.

[36] J. Bols, J. Degroote, B. Trachet, B. Verhegghe, P. Segers, and J. Vierendeels, “A computational method to assess the in vivo stresses and unloaded configuration of patient-specific blood vessels,” Journal of Computational and Applied Mathematics, vol. 246, pp. 10–17, 2013/07/01/ 2013, doi: https://doi.org/10.1016/j.cam.2012.10.034.

[37] M. Liu et al., “Computation of a probabilistic and anisotropic failure metric on the aortic wall using a machine learning-based surrogate model,” Computers in Biology and Medicine, vol. 137, p. 104794, 2021/10/01/ 2021, doi: https://doi.org/10.1016/j.compbiomed.2021.104794.

[38] K. Miller and J. Lu, “On the prospect of patient-specific biomechanics without patient-specific properties of tissues,” Journal of the Mechanical Behavior of Biomedical Materials, vol. 27, pp. 154–166, 2013/11/01/ 2013, doi: https://doi.org/10.1016/j.jmbbm.2013.01.013.

[39] G. R. Joldes, K. Miller, A. Wittek, and B. Doyle, “A simple, effective and clinically applicable method to compute abdominal aortic aneurysm wall stress,” Journal of the Mechanical Behavior of Biomedical Materials, vol. 58, pp. 139–148, 2016/05/01/ 2016, doi: https://doi.org/10.1016/j.jmbbm.2015.07.029.

[40] J. D. Humphrey and S. K. Kyriacou, “The use of Laplace’s equation in aneurysm mechanics,” Neurological Research, vol. 18, no. 3, pp. 204–208, 1996/06/01 1996, doi: 10.1080/01616412.1996.11740404.

[41] M. Liu, L. Liang, and W. Sun, “A new inverse method for estimation of in vivo mechanical properties of the aortic wall,” Journal of the Mechanical Behavior of Biomedical Materials, vol. 72, pp. 148–158, 8// 2017, doi: https://doi.org/10.1016/j.jmbbm.2017.05.001.

[42] L. Pineda et al., “Theseus: A Library for Differentiable Nonlinear Optimization,” Advances in Neural Information Processing Systems, 2022.

[43] I. J. Goodfellow et al., “Generative Adversarial Nets,” Advances in Neural Information Processing Systems, pp. 2672–2680, 2014.

[44] J. Sohl-Dickstein, E. A. Weiss, N. Maheswaranathan, and S. Ganguli, “Deep unsupervised learning using nonequilibrium thermodynamics,” presented at the Proceedings of the 32nd International Conference on International Conference on Machine Learning-Volume 37, Lille, France, 2015.

[45] M. Liu, L. Liang, and W. Sun, “A generic physics-informed neural network-based constitutive model for soft biological tissues,” Computer Methods in Applied Mechanics and Engineering, vol. 372, p. 113402, 2020/12/01/ 2020, doi: https://doi.org/10.1016/j.cma.2020.113402.

[46] V. Tac, V. D. Sree, M. K. Rausch, and A. B. Tepole, “Data-driven modeling of the mechanical behavior of anisotropic soft biological tissue,” Engineering with Computers, vol. 38, no. 5, pp. 4167–4182, 2022/10/01 2022, doi: 10.1007/s00366-022-01733-3.

[47] P. Chen and J. Guilleminot, “Polyconvex neural networks for hyperelastic constitutive models: A rectification approach,” Mechanics Research Communications, vol. 125, p. 103993, 2022/10/01/ 2022, doi: https://doi.org/10.1016/j.mechrescom.2022.103993.

[48] V. Tac, F. Sahli Costabal, and A. B. Tepole, “Data-driven tissue mechanics with polyconvex neural ordinary differential equations,” Computer Methods in Applied Mechanics and Engineering, vol. 398, p. 115248, 2022/08/01/ 2022, doi: https://doi.org/10.1016/j.cma.2022.115248.

[49] Y. Leng, V. Tac, S. Calve, and A. B. Tepole, “Predicting the mechanical properties of biopolymer gels using neural networks trained on discrete fiber network data,” Computer Methods in Applied Mechanics and Engineering, vol. 387, p. 114160, 2021/12/15/ 2021, doi: https://doi.org/10.1016/j.cma.2021.114160.

[50] T. Gärtner, M. Fernández, and O. Weeger, “Nonlinear multiscale simulation of elastic beam lattices with anisotropic homogenized constitutive models based on artificial neural networks,” Computational Mechanics, vol. 68, no. 5, pp. 1111–1130, 2021/11/01 2021, doi: 10.1007/s00466-021-02061-x.

[51] Z. Jiang, J. Choi, and S. Baek, “Machine learning approaches to surrogate multifidelity Growth and Remodeling models for efficient abdominal aortic aneurysmal applications,” Computers in Biology and Medicine, vol. 133, p. 104394, 2021/06/01/ 2021, doi: https://doi.org/10.1016/j.compbiomed.2021.104394.

[52] M. Raissi, P. Perdikaris, and G. E. Karniadakis, “Physics-informed neural networks: A deep learning framework for solving forward and inverse problems involving nonlinear partial differential equations,” Journal of Computational Physics, vol. 378, pp. 686–707, 2019/02/01/ 2019, doi: https://doi.org/10.1016/j.jcp.2018.10.045.

[53] D. W. Abueidda, S. Koric, E. Guleryuz, and N. A. Sobh, “Enhanced physics-informed neural networks for hyperelasticity,” International Journal for Numerical Methods in Engineering, vol. 124, no. 7, pp. 1585–1601, 2023/04/15 2023, doi: https://doi.org/10.1002/nme.7176.

[54] J. Wang, C. Wang, Q. Lin, C. Luo, C. Wu, and J. Li, “Adversarial attacks and defenses in deep learning for image recognition: A survey,” Neurocomputing, vol. 514, pp. 162–181, 2022/12/01/ 2022, doi: https://doi.org/10.1016/j.neucom.2022.09.004.

[55] A.-K. Dombrowski, M. Alber, C. J. Anders, M. Ackermann, K.-R. Müller, and P. Kessel, “Explanations can be manipulated and geometry is to blame,” in Proceedings of the 33rd International Conference on Neural Information Processing Systems: Curran Associates Inc., 2019, p. Article 1217.

[56] J. Heo, S. Joo, and T. Moon, “Fooling neural network interpretations via adversarial model manipulation,” in Proceedings of the 33rd International Conference on Neural Information Processing Systems: Curran Associates Inc., 2019, p. Article 263.

[57] S. A. Maas, B. J. Ellis, G. A. Ateshian, and J. A. Weiss, “FEBio: finite elements for biomechanics,” (in eng), J Biomech Eng, vol. 134, no. 1, p. 011005, Jan 2012, doi: 10.1115/1.4005694.

[58] O. C. Zienkiewicz and R. L. Taylor, The Finite Element Method: Its Basis and Fundamentals. Elsevier, 2013.

[59] A. Logg, K.-A. Mardal, and G. Wells, Automated Solution of Differential Equations by the Finite Element Method: The FEniCS Book. Springer Berlin, Heidelberg, 2012.

[60] A. Elouneg, D. Sutula, J. Chambert, A. Lejeune, S. P. A. Bordas, and E. Jacquet, “An open-source FEniCS-based framework for hyperelastic parameter estimation from noisy full-field data: Application to heterogeneous soft tissues,” Computers & Structures, vol. 255, p. 106620, 2021/10/15/ 2021, doi: https://doi.org/10.1016/j.compstruc.2021.106620.

[61] S. K. Mitusch, S. W. Funke, and M. Kuchta, “Hybrid FEM-NN models: Combining artificial neural networks with the finite element method,” Journal of Computational Physics, vol. 446, p. 110651, 2021/12/01/ 2021, doi: https://doi.org/10.1016/j.jcp.2021.110651.

